# BRAVE: a highly accurate method for predicting HIV-1 antibody resistance using large language models for proteins

**DOI:** 10.1101/2025.07.28.667234

**Authors:** Mohammed El Anbari, Tatsiana Bylund, Sijy O’Dell, Emily Tourtellott, Krisha McKee, Stephen D Schmidt, Nonhlanhla N. Mkhize, Penny L Moore, Nicole Doria-Rose, Tongqing Zhou, Reda Rawi

## Abstract

**Motivation:** Broadly neutralizing antibodies (bNAbs) that target the envelope glycoprotein (Env) of human immunodeficiency virus-1 (HIV-1) have been utilized in clinical trials aimed at preventing and treating HIV-1 infections. However, the emergence of neutralization resistance to bNAbs occurs rapidly due to the high mutation rate of HIV-1. Previous studies have suggested the use of *in silico* methods to effectively predict the resistance of HIV-1 isolates to bNAbs. In this study, we present a novel machine learning approach called BRAVE (Bnab Resistance Analysis Via Evolutionary scale modeling 2) designed to predict HIV-1 resistance against 33 known bNAbs. This innovative tool employs a Random Forests classifier that uses a protein language model to reliably capture protein features.

**Results:** BRAVE outperformed leading resistance prediction tools on various performance metrics, attaining the highest performance in established classification measures including accuracy, area under the curve, logarithmic loss, and F1-score. Importantly, rigorous statistical comparisons (p<0.001) show that BRAVE is significantly more accurate than state-of-the-art neutralization prediction tools. BRAVE will facilitate informed decisions of antibody usage and sequence-based monitoring of viral escape in clinical settings.

**Availability and implementation:** BRAVE software is available for download under GitHub (https://github.com/kiryst/BRAVE/tree/master).

**Supplementary information:** Supplementary data are available at Bioinformatics online.

## 1 Introduction

HIV-1 bNAbs neutralize a wide range of HIV-1 strains with some showing extraordinary potency. Numerous of these bNAbs have demonstrated protection from viral challenges in animal models even at low concentrations in serum (Gautam, et al., 2016; Moldt, et al., 2016; Moldt, et al., 2012; Pegu, et al., 2014; Rudicell, et al., 2014; Saunders, et al., 2015; Shingai, et al., 2014), motivating the application of bNAbs for prevention and treatment in human populations. However, HIV-1’s exceptionally high mutation rate enables the virus to develop resistance to bNAbs (Yu, et al., 2019). Hence, the administration of bNAbs, even to strains that are initially sensitive, can result in viral escape, diminishing or completely negating the effectiveness of bNAbs (Caskey, et al., 2017). Therefore, an accurate tool for predicting resistance to HIV-1 antibodies would be highly beneficial in selecting the appropriate antibody for administration and in tracking viral escape throughout the treatment process. Additionally, given the ongoing clinical trials involving various bNAbs, there is an urgent requirement for *in silico* pipeline able to accurately assess the antibody resistance identified in these trials. Commonly, antibody neutralization capacity is assessed using the virus pseudotyped TZM-bl neutralization assay which is laborious and cost-intensive. Multiple *in silico* tools to tackle this challenge have been proposed, utilizing various machine learning (ML) techniques. Bayesian graphical models have been employed in (Hepler, et al., 2014), artificial feedforward neural networks in (Buiu, et al., 2016), non-linear support vector machines with a string kernel in (Hake and Pfeifer, 2017), and gradient boosting machines in (Rawi, et al., 2019). Atomistic modeling combined with ML has also been explored in (Conti and Karplus, 2019), along with Bayesian ML models (Yu, et al., 2019), and nonparametric ensemble-based cross-validated learning methods (Magaret, et al., 2019). Additionally, neural networks and transfer learning integrating both virus and antibody sequences have been applied in (Danaila and Buiu, 2022), as well as multi-task learning approaches (Igiraneza, et al., 2024). In this study, we introduce BRAVE, a ML tool that predicts bNAb neutralization resistance leveraging the recent advances in protein language models (PLMs). BRAVE combines the strengths of Evolutionary scale modeling-2 (ESM-2) (Lin, et al., 2023), a transformer-based PLM trained on massive protein sequence data, which provides meaningful representations of amino acid sequences, with a non-linear Random Forests (RF) classifier to accurately predict neutralization resistance for 33 bNAbs. Rigorous statistical comparisons shows that BRAVE significantly outperformed (p<0.001) state-of-the-art bNAb resistance prediction tools in terms of accuracy, area under the curve, logarithmic loss, and F1-score, showing its overall superiority. In addition, BRAVE is proving its generalizability by performing best on independent data sets that were not used during model training.

## 2 Methods

### 2.1 Algorithm

Figure 1 outlines the BRAVE pipeline flowchart. First, HIV-1 sequences are aligned using MAFFT sequence alignment tool. Next, sequence embeddings are generated with the ESM-2 protein language model. IC_50_ values are binarized using a 50□µg/mL threshold, and a RF classifier is trained on the resulting dataset. Model performance is evaluated using five-fold nested cross-validation and assessed on both training and independent test sets using standard metrics.

**Figure 1.**
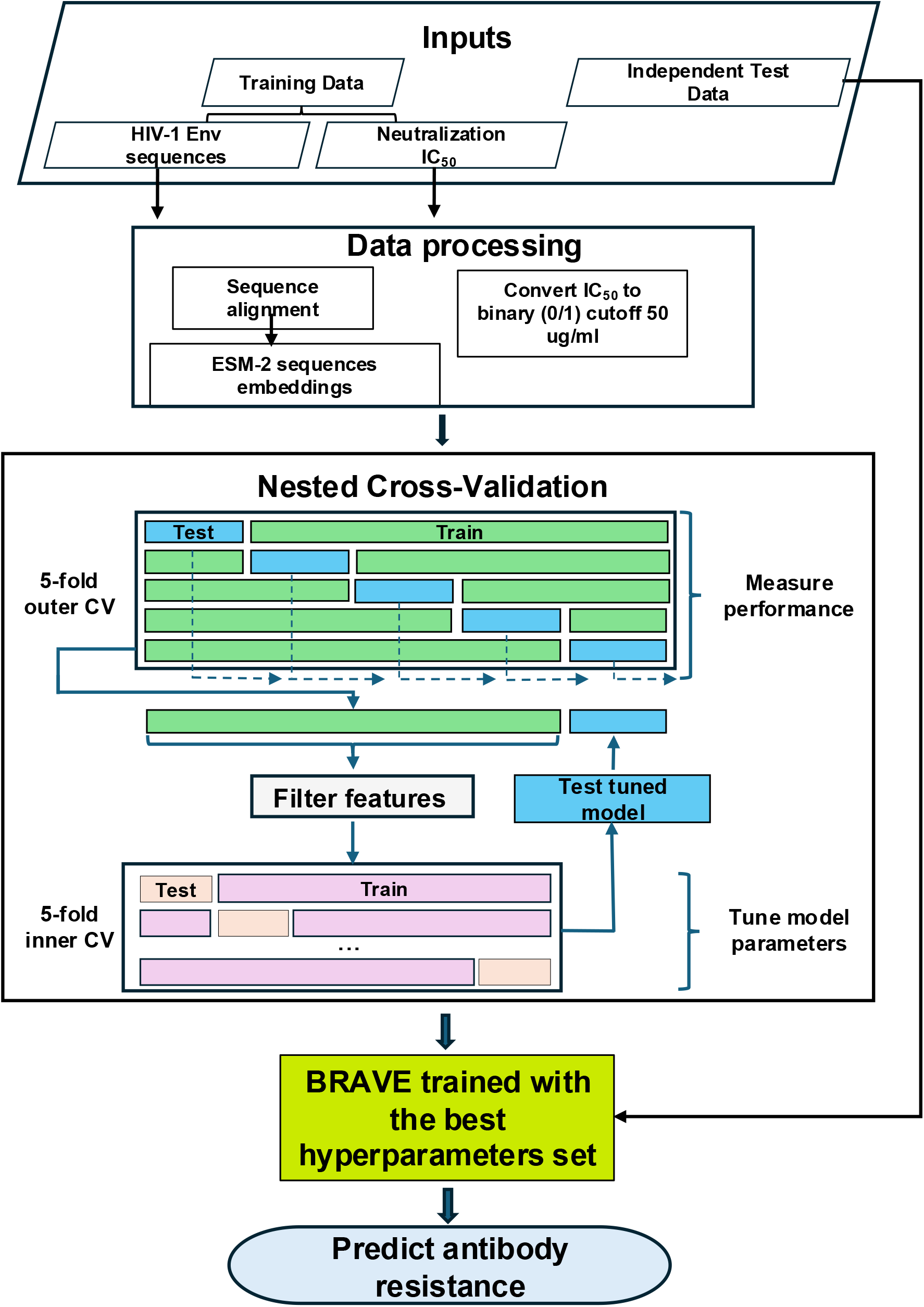
Flowchart of the BRAVE pipeline for predicting antibody resistance.

### 2.2 Data preprocessing

#### 2.2.1 Training Datasets

BRAVE was trained on sequence and neutralization data for 33 HIV-1 bNAbs. All datasets were preprocessed in a recent study (Igiraneza et al., 2024) and are publicly available via GitHub: https://github.com/iaime/LBUM. The data originate from the CATNAP database (Yoon et al., 2015). Left-censored IC_50_ neutralization values (IC_50_ < DL, with DL=detection limit) were imputed using the DL. Since no right-censored values were present, the geometric mean was used to aggregate IC50 values for Env sequences with multiple measurements. The IC50 outcome was binarized: sequences with IC_50_ ≥ 50□µg/mL or right-censored values were labeled as resistant; those with IC_50_ < 50□µg/mL were labeled as sensitive.

As shown in Figure S1, the 33 bNAb datasets vary in size (183 to 843 sequences) and class imbalance, with majority-to-minority class ratios ranging from 1.02 to 10.21, reflecting heterogeneity in resistance profiles.

#### 2.2.2 Testing Datasets

To evaluate BRAVE’s performance on independent data, we assessed HIV-1 neutralization resistance against nine antibodies using test datasets from randomized clinical trials (Corey et al., 2021; Mkhize et al., 2023). These datasets were not included in the training phase and provide an external benchmark for model generalizability. Details on sample size and class distribution are shown in Figure S2. Sample sizes range from 15 to 56, with class imbalance ratios between 1.28 and 3.00.

#### 2.2.3 Sequence identity

To assess sequence relatedness, pairwise sequence identity was computed for both training and test datasets. In the training set, sequence identity ranges from 56% to 99.99%, reflecting broad sequence diversity. The test set shows less variability, with identity values ranging from 74% to 99.99%. Within the training set, identity between resistant and sensitive sequences spans 65% to 99.99%, with differences as small as a single amino acid substitution. This reflects the potential for a single residue change— particularly at the antibody interface to enable resistance. In contrast, the test set exhibits a narrower identity range between resistant and sensitive sequences, from 79% to 95%.

### 2.3 Feature engineering using protein language models

PLMs have demonstrated strong performance in protein engineering and function prediction tasks (Alley et al., 2019; Meier et al., 2021; Rives et al., 2021). PLMs apply techniques from natural language processing to protein sequences by encoding each sequence into a numerical vector, or embedding, designed to capture biochemical and evolutionary properties. Similar sequences tend to yield similar embeddings, facilitating downstream learning tasks. In this study, we used the ESM-2 family of PLMs (Igiraneza et al., 2023), which were trained on hundreds of millions of protein sequences, enhancing their ability to model sequence relationships. To ensure uniform embedding dimensions, all sequences were aligned using MAFFT (Katoh et al., 2002).

### 2.4 Random Forests

To build the training models, we used RF, a method introduced by Leo Breiman (Breiman, 2001). RF is a nonparametric approach widely applied across various fields (Biau and Scornet, 2016). It is well-known for its excellent predictive performance and versatility. It can handle different types of data and is particularly effective for high-dimensional datasets, where the number of features exceeds largely the sample size, as well as for capturing complex, non-linear relationships between the outcome and the features. In this work, we used the R package ranger (Wright and Ziegler, 2017), which is a fast implementation of RF. We performed a grid search to optimize the hyperparameters, including the number of trees in the RF (*mtry*), the minimum number of observations required in a node (*min*.*node*.*size*), and the splitting criterion used to determine the best split at each node (*splitrule*). We considered a broad grid of values for *mtry* consisting of small and large values and depending on the number of features (#features=100,000). We created a grid consisting of *mtry* × *min*.*node*.*size* × *splitrule* hyperparameters, where

- mtry ∈ {2, 4, 6, 8, 10, log10(#features), 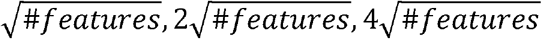, #features * 0.01, #features * 0.05, #features * 0.1},
- min. node. size ∈ {1, 2, 3, 4, 5, 6, 7, 8, 9, 10},
- splitrule ∈ {variance, extratrees}, resulting in 240 different combinations of hyperparameters. The optimal hyperparameters were chosen using nested cross-validation.

### 2.5 Nested cross-validation

In this work, we used the R package nestedcv to apply 5×5 nested cross-validation to estimate the performance of each model. The data was partitioned into outer and inner folds, as shown in Figure 1. The inner fold cross-validation is used to find the optimal hyperparameters of the model. The model is then fitted on the entire inner fold and tested on the left-out data from the outer fold. This process is repeated across all five outer folds, and predictions along with ground truth label vectors from the unseen data in the outer folds are used to calculate various performance metrics. The average performance over the five outer folds is used to compare different neutralization prediction techniques.

### 2.6 Compared methods and metrics for classifier evaluation

The performance of the method BRAVE was compared to two state-of-the-art antibody neutralization predictors: bNAb-Resistance Predictor (bNAb-ReP), which uses a gradient boosting machine, and Language-based universal model (LBUM), which is based on a two-layer bidirectional Long Short-Term Memory. We compared the methods using various metrics, including accuracy (ACC), Area Under the Curve (AUC), Logarithmic Loss (Log-Loss), and F1-score (F1-Score). Each metric captures a different aspect of performance. ACC measures the proportion of correctly classified instances out of the total instances. AUC is interpreted as the probability that a model ranks resistant sequences higher than sensitive ones. The Log-Loss values are showing how close the predicted probabilities to the actual values. A lower Log-Loss value indicates better model performance. The F1-score which is a combination of the information from precision and recall into a single metric by calculating their harmonic mean. This metric is particularly useful when dealing with imbalanced datasets.

### 2.7 Robust statistical comparisons of antibody neutralization predictors

We used robust statistical methods, as proposed in (Bergmann and Hommel, 1988; Demšar, 2006), to compare the performance of the different antibody neutralization predictors across 33 bNAb datasets. These papers have significantly influenced ML research by promoting robust statistical methodologies for evaluating classifier performance over multiple data sets. To reduce sensitivity to outliers, the authors propose to compare the classifiers based on the average ranks of the different performance measures, rather than the raw metrics themselves. A Friedman test is first used on the data of ranks. If the Friedman’s test identifies significant differences, a post-hoc Nemenyi test is performed to compare all pairs of classifiers. Finally, a Bergmann and Hommel dynamic correction is used for multiple hypotheses testing.

## 3 Results

### 3.1 BRAVE training

BRAVE was developed utilizing sequence and neutralization data for 33 bNAbs against HIV-1, sourced from the CATNAP database. The training of classifiers involved two primary phases: the generation of features and the training of a RF classifier (Fig. 1). ESM-2 embeddings were employed to present full-length HIV-1 Env sequences as features (see Methods), and the training of the RF model included a hyperparameter optimization process to identify the optimal parameter set for each bNAb-related classifier (see Methods).

Next, we evaluated the training performance of all 33 BRAVE classifiers compared to the state-of-the-art models LBUM and bNAb-ReP using five-fold nested cross-validation (see Methods). BRAVE outperformed LBUM and bNAb-ReP in accuracy, being superior in for 25 bNAb across epitope categories (6 for LBUM and 3 for bNAb-ReP) (Table 1). The prediction performance of BRAVE was also the best in terms of other performance metrics, such as AUC (Table 1), Log-Loss, F1-score (both Table 2), and others (Table S3).

**Table 1.**
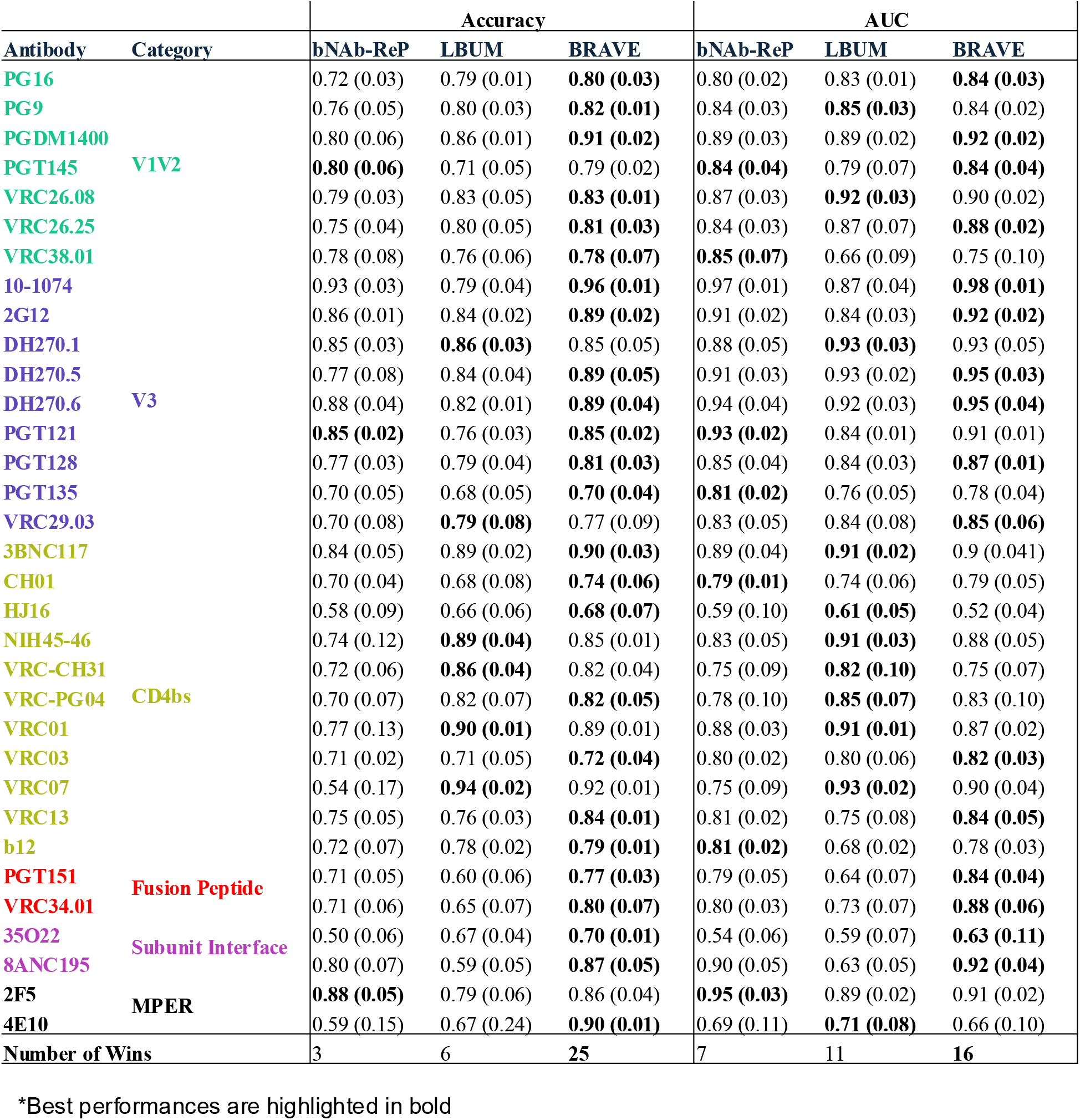
Accuracy and AUC of BRAVE compared to state-of-the-art antibody neutralization predictors.* *Best performances are highlighted in bold

**Table 2.**
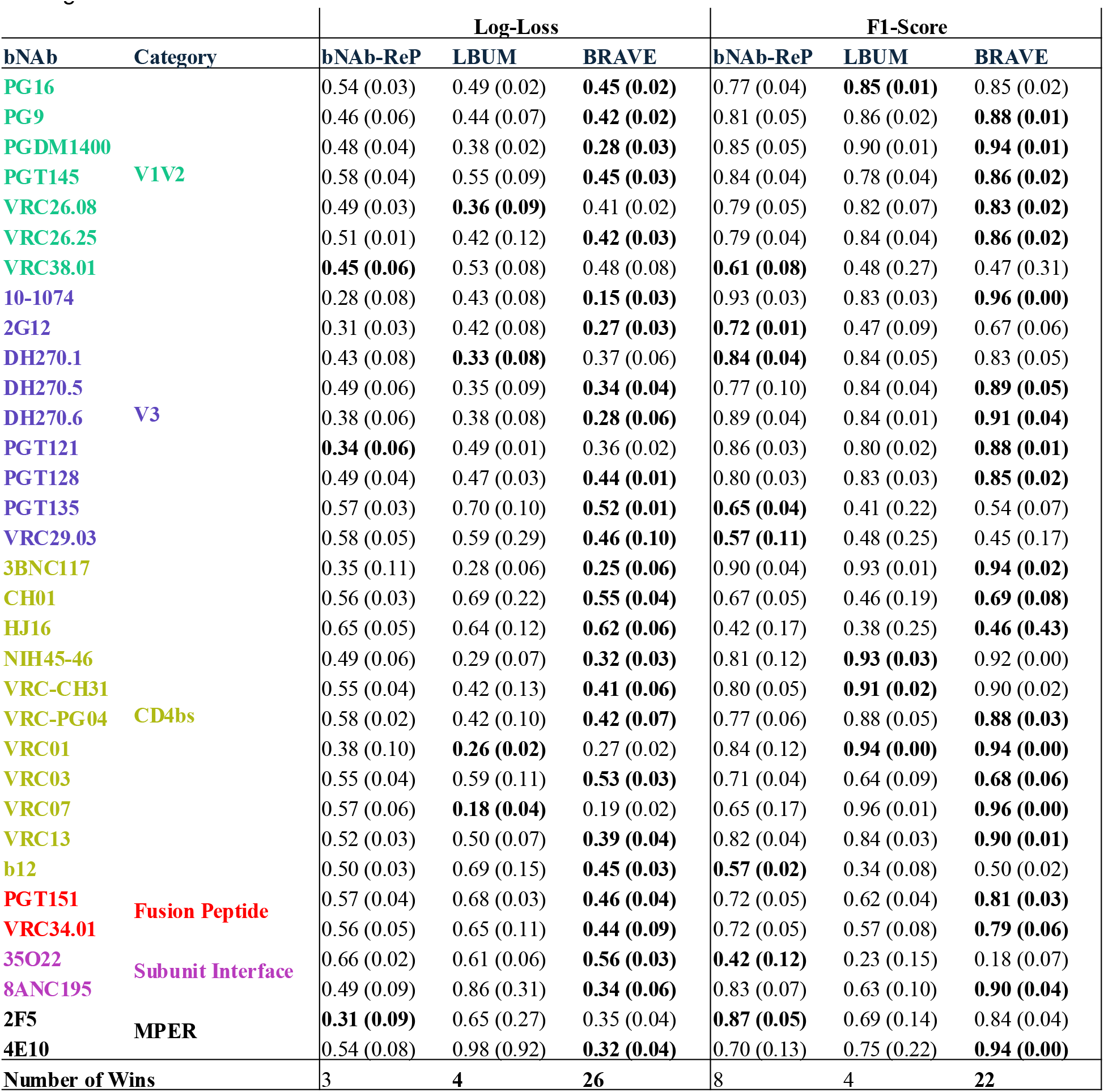
Comparison of BRAVE’s performance with leading antibody neutralization predictors in terms of log loss and F1-Score.

Next, we applied the Nemenyi post-hoc test in conjunction with a Friedman significance test to assess the statistical differences among the methods employed. BRAVE demonstrated a significantly higher accuracy compared to both bNAb-ReP and LBUM (p < 0.001), whereas no significant difference was found between bNAb-ReP and LBUM (p = 0.0849) in terms of accuracy (Figure 2). Regarding the AUC performance metric, BRAVE was significantly more accurate than LBUM (p = 0.0101) but did not show a significant difference when compared to bNAb-ReP (p = 0.0785). Additionally, there was no significant difference in AUC between bNAb-ReP and LBUM (p = 0.2403). Similar patterns were observed for the performance metrics of Log-Loss and F1-score, with BRAVE significantly outperforming both bNAb-ReP and LBUM. These results underscore the exceptional performance of BRAVE across the four assessed performance metrics.

**Figure 2.**
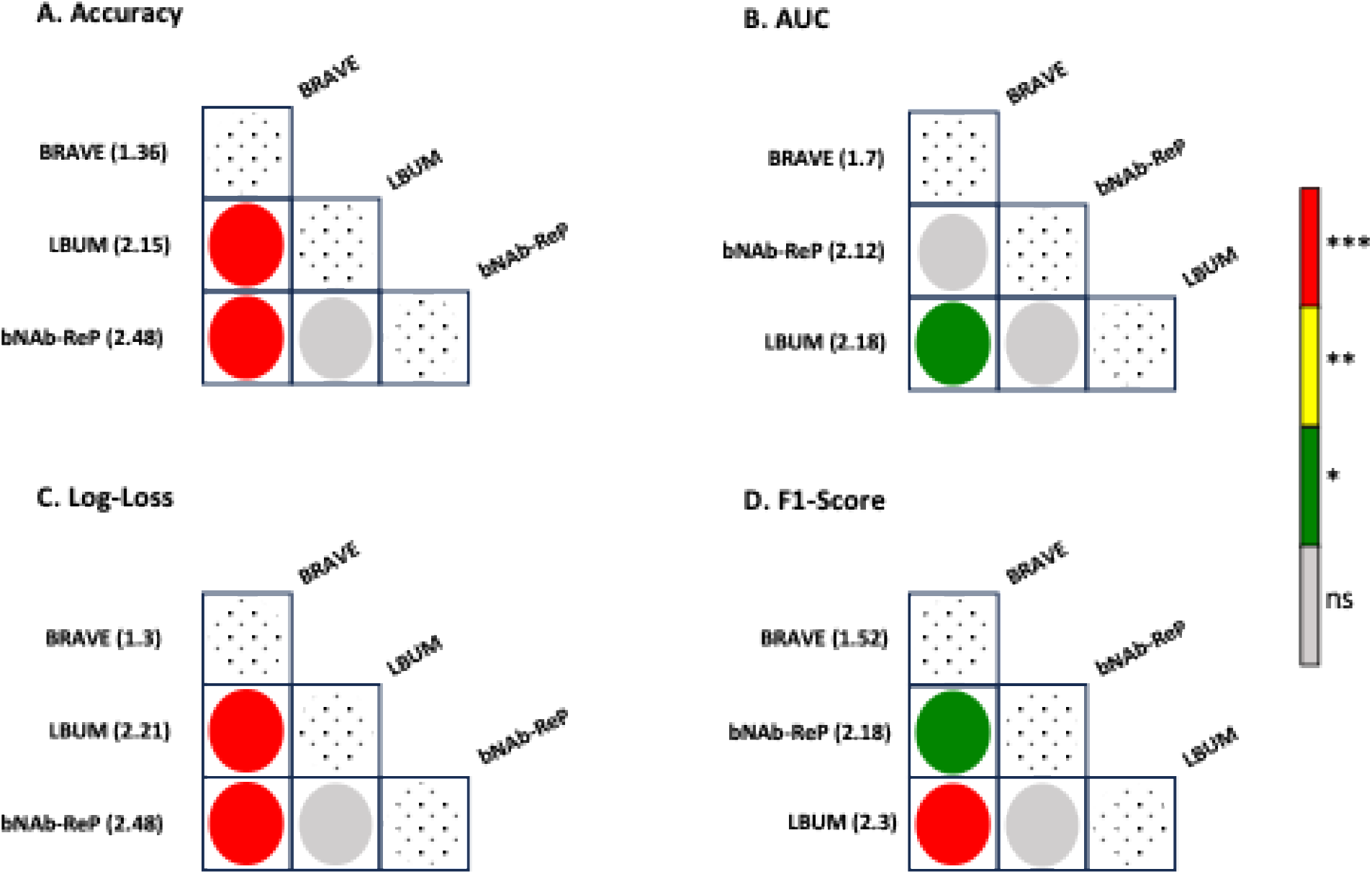
BRAVE significantly outperforms state-of-the-art neutralization predictors. Statistical comparisons of bNAb-ReP, LBUM, and BRAVE across 33 bNAbs show that BRAVE significantly outperforms other state-of-the-art neutralization predictors in terms of accuracy, AUC, log loss, and F1-Score. Adjusted p-values less than 0.05 were considered significant (* p < 0.05; ** p < 0.01; *** p < 0.001).

### 3.2 Antibody resistance prediction on independent test set

To assess the efficacy of BRAVE beyond the training datasets derived from CATNAP, we evaluated the neutralization resistance of HIV-1 isolates in response to nine different antibodies. The test data sets that we are using were provided in (Corey, et al., 2021) and (Mkhize, et al., 2023).

The performance of BRAVE demonstrated an advantage over bNAb-ReP when analyzed through four key performance metrics: accuracy, AUC, Log-Loss, and F1-score. Specifically, BRAVE achieved superior results in six out of ten instances for accuracy, six out of nine for AUC, five out of nine for Log-Loss, and five out of nine for F1-score (Table 3). BRAVE achieves an accuracy between 0.56 and 0.94 (median: 0.80), compared to 0.46–0.95 for bNAb-ReP (median: 0.73). With respect to AUC, BRAVE ranges from 0.55 to 0.92 (median: 0.85), while bNAb-ReP ranges from 0.59 to 0.89 (median: 0.74). Log-Loss values span 0.29-0.85 for BRAVE and 0.29-0.73 for bNAb-ReP, with a lower median for BRAVE (0.44 vs. 0.54). The F1-score ranges from 0.36 to 0.90 for BRAVE (median: 0.72) and from 0.40 to 0.89 for bNAb-ReP (median: 0.73).

**Table 3.**
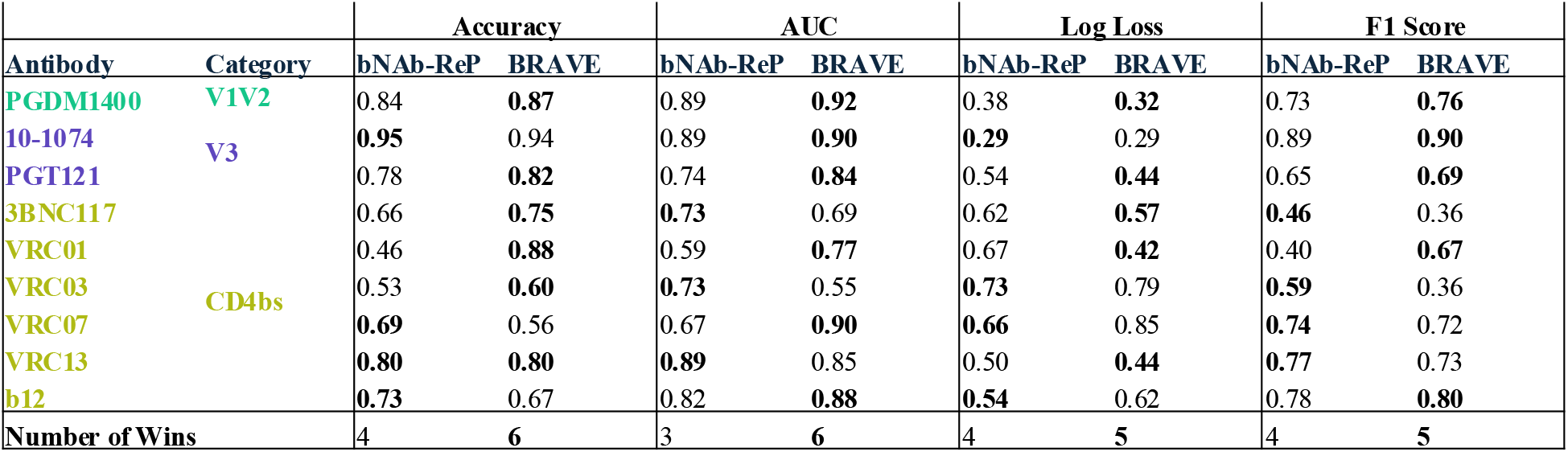
BRAVE achieves higher success rate compared to bNAb-ReP on independent test datasets.

## Conclusion

The advancement of accurate *in silico* sequence-based predictions for bNAb neutralization resistance remains crucial in the domain of HIV-1 research. In this study, we present BRAVE, an innovative ML framework designed to predict HIV-1 resistance to 33 bNAbs. This approach integrates the established predictive capabilities of a RF classifier with feature engineering utilizing ESM-2 protein language models. Nested cross-validation was employed to ensure that all evaluations were performed on data partitions not used during model training, thereby preventing information leakage and enabling an unbiased assessment of model performance. Statistical analyses demonstrate that BRAVE outperforms current leading prediction tools across various performance metrics, including accuracy, AUC, Log-Loss, and F1-Score. Furthermore, the proposed method exhibits generalizability by effectively predicting resistance in an independent test dataset. We note here that the results of LBUM are not reported because the authors do not provide a final model trained on the entire dataset that can be used to predict resistance on independent test data. Instead, their code provides results on five splits of the training data, and the resulting performance on the independent test set was notably low.

A potential direction for future work is the prediction of resistance to a combination of monoclonal bNAbs via multi-task learning. Additionally, interpretability of ESM-2 embedding is challenging due to their high dimensionality and the complex non-linear transformations used to generate them, which obscure the relationships between individual features and the embeddings. Hence another potential enhancement could be achieved by applying feature selection methods to ESM-type embeddings, which would facilitate a deeper comprehension of the mechanisms and enable the extraction of features to gain insights into critical elements that dictate neutralization resistance.

## Acknowledgements

We thank J. Stuckey for assistance with the figures. We thank all members of SBS and SBIC VRC NIAID for their feedback. This work utilized the computational resources of the NIH HPC Biowulf cluster (NIH HPC). This study used the Office of Cyber Infrastructure and Computational Biology (OCICB) High Performance Computing (HPC) cluster at the National Institute of Allergy and Infectious Diseases (NIAID), Bethesda, MD.

---

**Figure S1.**
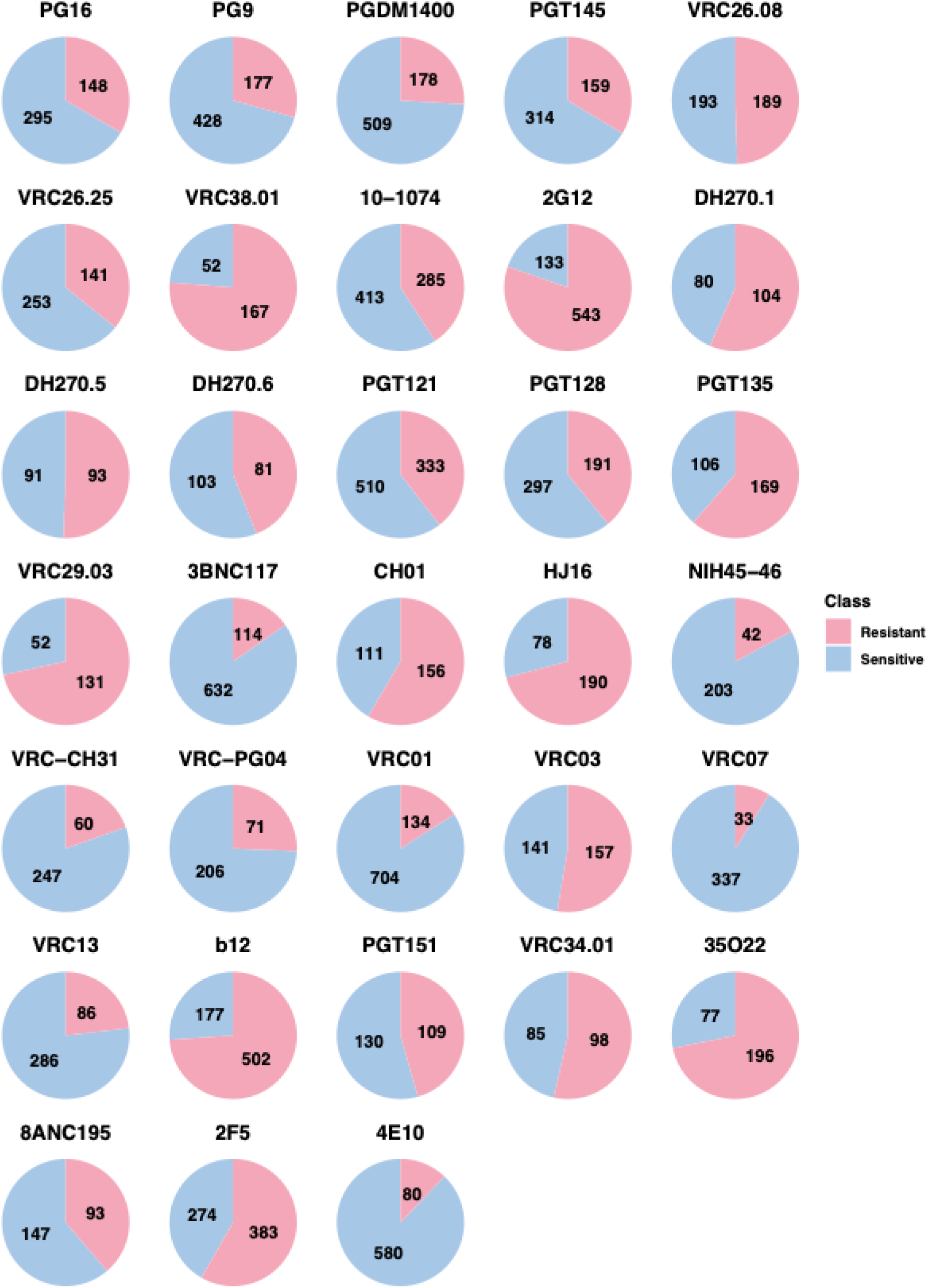
Training sets class distributions. Pie charts showing the distribution of “Resistant” and “Sensitive” classes for each bNAb training data, highlighting different imbalance ratios.

**Figure S2.**
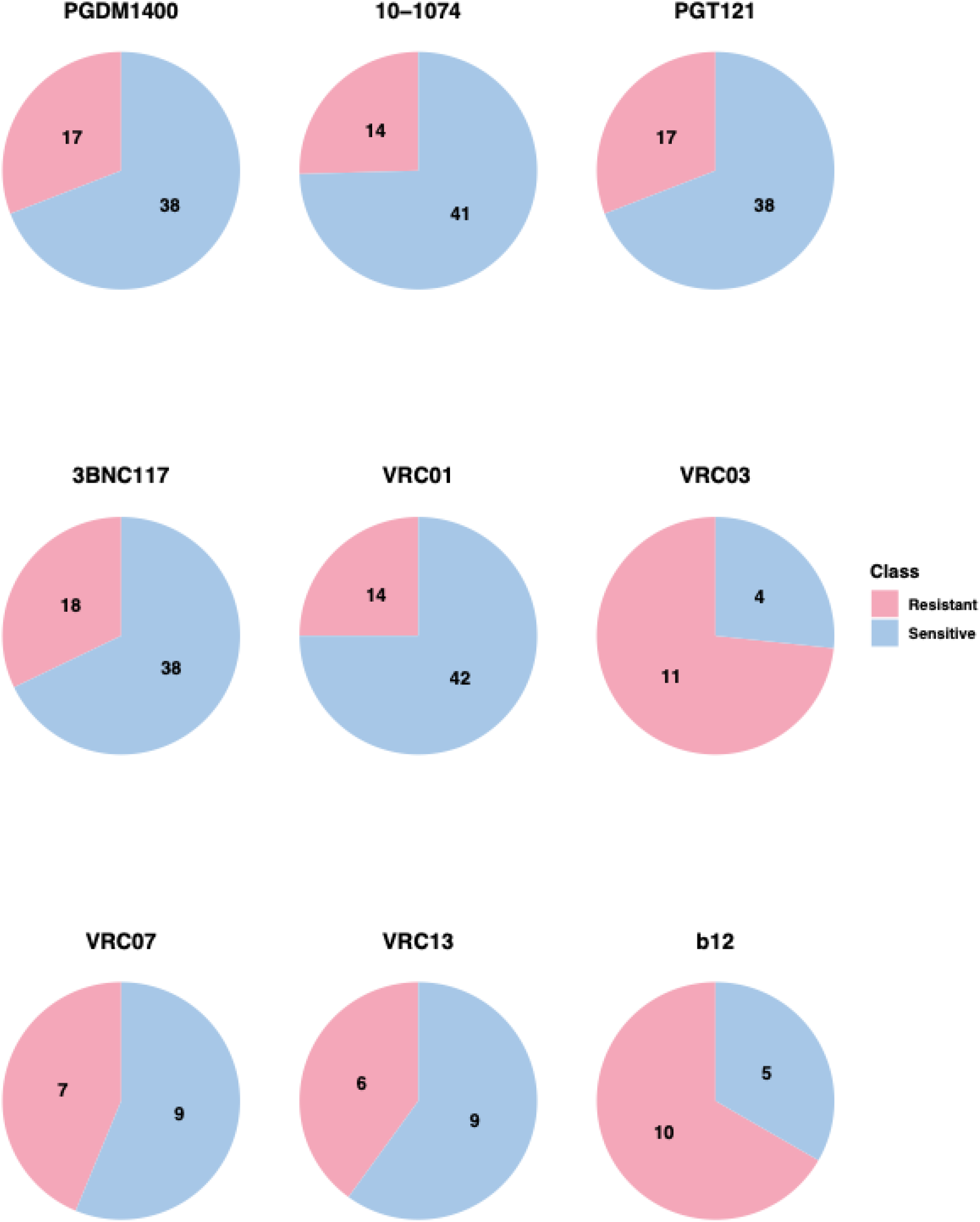
Test sets class distributions Pie charts showing the distribution of “Resistant” and “Sensitive” classes for each bNAb test data, highlighting different imbalance ratios.

## References

Alley, E.C., et al. Unified rational protein engineering with sequence-based deep representation learning. Nat Methods 2019;16(12):1315–1315.

Bergmann, B. and Hommel, G. Improvements of general multiple test procedures for redundant systems of hypotheses. Multiple hypotheses testing 1988;70:100-115.

Biau, G. and Scornet, E. A random forest guided tour. Test-Spain 2016;25(2):197–197.

Breiman, L. Random forests. Mach Learn 2001;45(1):5–5.

Buiu, C., Putz, M.V. and Avram, S. Learning the Relationship between the Primary Structure of HIV Envelope Glycoproteins and Neutralization Activity of Particular Antibodies by Using Artificial Neural Networks. Int J Mol Sci 2016;17(10).

Caskey, M., et al. Antibody 10-1074 suppresses viremia in HIV-1-infected individuals. Nat Med 2017;23(2):185–185.

Conti, S. and Karplus, M. Estimation of the breadth of CD4bs targeting HIV antibodies by molecular modeling and machine learning. Plos Computational Biology 2019;15(4).

Corey, L., et al. Two Randomized Trials of Neutralizing Antibodies to Prevent HIV-1 Acquisition. N Engl J Med 2021;384(11):1003–1003.

Danaila, V.R. and Buiu, C. Prediction of HIV sensitivity to monoclonal antibodies using aminoacid sequences and deep learning. Bioinformatics 2022;38(18):4278–4278.

Demšar, J. Statistical comparisons of classifiers over multiple data sets. Journal of Machine learning research 2006;7(Jan):1–30.

Gautam, R., et al. A single injection of anti-HIV-1 antibodies protects against repeated SHIV challenges. Nature 2016;533(7601):105–105.

Hake, A. and Pfeifer, N. Prediction of HIV-1 sensitivity to broadly neutralizing antibodies shows a trend towards resistance over time. Plos Computational Biology 2017;13(10).

Hepler, N.L., et al. IDEPI: rapid prediction of HIV-1 antibody epitopes and other phenotypic features from sequence data using a flexible machine learning platform. PLoS Comput Biol 2014;10(9):e1003842.

Igiraneza, A.B., et al. Learning patterns of HIV-1 resistance to broadly neutralizing antibodies with reduced subtype bias using multi-task learning. PLoS Comput Biol 2024;20(11):e1012618.

Katoh, K., et al. MAFFT: a novel method for rapid multiple sequence alignment based on fast Fourier transform. Nucleic Acids Research 2002;30(14):3059–3059.

Lewis, M.J., et al. nestedcv: an R package for fast implementation of nested cross-validation with embedded feature selection designed for transcriptomics and high-dimensional data. Bioinformatics Advances 2023;3(1).

Lin, Z., et al. Evolutionary-scale prediction of atomic-level protein structure with a language model. Science 2023;379(6637):1123–1123.

Magaret, C.A., et al. Prediction of VRC01 neutralization sensitivity by HIV-1 gp160 sequence features. Plos Computational Biology 2019;15(4).

Meier, J., et al. Language models enable zero-shot prediction of the effects of mutations on protein function. Adv Neur In 2021;34.

Mkhize, N.N., et al. Neutralization profiles of HIV-1 viruses from the VRC01 Antibody Mediated Prevention (AMP) trials. PLoS Pathog 2023;19(6):e1011469.

Moldt, B., et al. Neutralizing antibody affords comparable protection against vaginal and rectal simian/human immunodeficiency virus challenge in macaques. AIDS 2016;30(10):1543–1543.

Moldt, B., et al. Highly potent HIV-specific antibody neutralization in vitro translates into effective protection against mucosal SHIV challenge in vivo. Proc Natl Acad Sci U S A 2012;109(46):18921–18921.

Pegu, A., et al. Neutralizing antibodies to HIV-1 envelope protect more effectively in vivo than those to the CD4 receptor. Sci Transl Med 2014;6(243).

Rawi, R., et al. Accurate Prediction for Antibody Resistance of Clinical HIV-1 Isolates. Sci Rep-Uk 2019;9.

Rives, A., et al. Biological structure and function emerge from scaling unsupervised learning to 250 million protein sequences. P Natl Acad Sci USA 2021;118(15).

Rudicell, R.S., et al. Enhanced potency of a broadly neutralizing HIV-1 antibody in vitro improves protection against lentiviral infection in vivo. J Virol 2014;88(21):12669–12669.

Saunders, K.O., et al. Sustained Delivery of a Broadly Neutralizing Antibody in Nonhuman Primates Confers Long-Term Protection against Simian/Human Immunodeficiency Virus Infection. J Virol 2015;89(11):5895–5895.

Shingai, M., et al. Passive transfer of modest titers of potent and broadly neutralizing anti-HIV monoclonal antibodies block SHIV infection in macaques. J Exp Med 2014;211(10):2061–2061.

Stone, M. An asymptotic equivalence of choice of model by cross-validation and Akaike’s criterion. Journal of the Royal Statistical Society: Series B (Methodological) 1977;39(1):44–44.

Wright, M.N. and Ziegler, A. ranger: A Fast Implementation of Random Forests for High Dimensional Data in C plus plus and R. J Stat Softw 2017;77(1):1–1.

Yoon, H., et al. CATNAP: a tool to compile, analyze and tally neutralizing antibody panels. Nucleic Acids Res 2015;43(W1):W213–219.

Yu, W.H., et al. Predicting the broadly neutralizing antibody susceptibility of the HIV reservoir. JCI Insight 2019;4(17).

